# Assessment of chemical methods in the removal of the spore coat and exosporium layers of *Clostridioides difficile* Spores

**DOI:** 10.1101/2024.04.07.588478

**Authors:** Javier Sanchez, Alba Romero-Rodriguez, Scarlett Troncoso-Cotal, Daniel Paredes-Sabja

## Abstract

*Clostridioides difficile* spores are essential for initiation, recurrence and transmission of the disease. The spore surface layers are composed of an outermost exosporium layer that surrounds another proteinaceous layer, the spore coat. These spore surfaces layers are responsible for initial interactions with the host and spore resistance properties contributing to transmission and recurrence of CDI. During spore-development, assembly of both layers is tightly interconnected thus studying the surface is essential for understanding the assembly of these layers and identification of potential targets for therapeutics. Several spore coat /exosporium extraction methods have utilized different extraction procedures making comparison across studies difficult and their impact on spore surface layer properties remains unclear. Here, we tested how commonly used chemical methods remove the spore coat /exosporium layers, analyzing treated-spores by phase contrast microscopy, transmission electron microscopy, western blotting, and lysozyme-triggered germination to functionally characterize the extraction efficiency of these treatment on these layers. Our results provide a systematic analysis and offer a platform for future spore coat and exosporium-related studies.

## Introduction

*Clostridioides difficile* (*C. difficile*) is a Gram-positive anaerobe that has become a leading cause of antibiotic associated diarrhea in developed countries^1, 2^. Treatment against *C. difficile* infections (CDI) resolve 95 % of primary cases; however, ∼15 – 30 % of recovered CDI patients develop subsequent recurrent episodes of CDI, which result in mortality rates increasing up to 30%^3^. Although two major clostridial toxins, TcdA and TcdB, are necessary for disease manifestation^4^, the production of *C. difficile* spores during infection is essential for recurrence^5, 6^.

*C. difficile* spores are metabolically dormant and naturally resistant to antibiotics, heat, ethanol, and various cleaning agents^6, 7^. *C. difficile* spores are made by a series of concentric layers that assemble under the control of the mother cell developmental program, specifically under the RNA polymerase sporulation-specific sigma factors, SigE and SigK ^6, 7^. The outermost exosporium layer of *C. difficile* spores play important roles in infection, recurrence and interaction with host molecules ^8–12^. Importantly, during late stages of sporulation development, *C. difficile* forms two distinctive spores that differ in the thickness of the exosporium morphotype, while the underlying spore coat remains identical ^6, 13, 14^.

Both, the spore coat and exosporium of *C. difficile* spores are proteinaceous layers made of > 50 and ∼ 10 - 20 proteins, respectively^15, 16^. Both layers reassemble a highly crosslinked protein matrix that can withstand various environmental insults^7^. Numerous studies on the spore coat and exosporium layer of *C. difficile* spores have varied in the methods to extraction of spore coat and exosporium layers ^10, 15–23^. Because these chemical methods dissociate the entire proteinaceous spore surface layers, they are thought to extract the spore coat and exosporium layers, at least in *C. difficile*^10, 15–23^. Early work utilized Lithium dodecyl sulfate sample buffer – 10% 2-mercaptoethanol followed by heating at 90°C for 10 min to remove the spore coat and exosporium layers of *C. diffiicle* 630 spores ^23^. Three additional methods have been utilized to remove these extracts. For example, follow-up work utilized 2 X Laemmli buffer (4% SDS, 10% 2-mercaptoethanol) followed by incubation for 10 min at 100 °C ^10, 15, 17^. Several proteomic studies used a lysis buffer with 6 M urea, 5 mM DTT in ammonium bicarbonate at pH 8.0 followed by bead beating at room temperature ^16, 21, 22^. Finally, a blend of 8 M urea, 2 M thiourea, 5% (w/v) SDS, 2% 2-mercaptoethanol, denominated EBB buffer, followed by incubation for 20 min at 95°C has also been used extensively mainly in sporulating cultures ^18–20^. Although different in their components, most of these approaches use: i) a chaotropic agent, urea or urea + thiourea to disrupt hydrogen bonds and results in the dissolution of hydrophobic residues within the protein^24^; ii) SDS to denature and solubilize proteins; iii) a reducing agent which can be 2-mercaptoethanol or dithiothreitol. Despite the well use of these different chemical extraction methods, it is unclear whether the spore coat and exosporium layer are efficiently removed through chemical methods, commonly employed in *C. difficile* spore-research.

Efficient fractionation of the spore surface layers is detrimental to understand the localization of spore surface constituents in each of these layers. In this regard, we demonstrated that enzymatic digestion with proteinase K or trypsin, or a mechanical treatment with sonication, efficiently removed only the exosporium layer of *C. difficile* spores ^15^. This allowed the identification of the exosporium proteome in spores of strain 630 ^15^; however, a similar validation with chemical extraction methods has not been conducted. Therefore, to improve our tool-box of methods to study *C. difficile* spore architecture and composition, in this work, in this work, provide a systematic analysis of how two chemical methods (i.e., USD and EBB); that are commonly employed to remove the spore coat /exosporium extracts for immunoblotting analysis, remove these layers and impact the functional aspects of *C. difficile* spores. In additional, Laemmli buffer was tested for its efficiency to extract alone when compared to USD and EBB. By utilizing two exosporium markers (i.e, CdeC and CdeM)^10, 25, 26^, and the the spore cortex lytic enzyme SleC as a marker for the peptidoglycan-spore coat interface ^27, 28^, we monitor the efficiency of extraction of these layers through immunoblotting and transmission electron microscopy.

## Materials and Methods

### Spore Preparation and Purification

*C. difficile* strain R20291 was grown in Brain Heart Infusion broth (BHIS) (Difco) supplemented with 0.5% yeast extract and 0.1% cysteine anaerobically at 37°C. Overnight cultures were grown in BHIS broth with 0.1% sodium taurocholate and 0.2% d-fructose. Cultures were diluted to an OD600 of 0.5 and 250 uL was plated on 70:30 sporulation media under anaerobic conditions for 5 days at 37°C^29^. Sporulating cultures were collected from plates, resuspended in ice-cold distilled water, and left in 4°C overnight to allow for lysis of vegetative cells. After incubation, cells were centrifuged and pellets washed with ice-cold distilled water 5 times followed by repeated centrifugation and resuspension to remove vegetative cell debris. Spores were separated using a 60% sucrose gradient and centrifuged for 20 min at 14,000 rpm, spores were then washed with ice-cold water to remove residual sucrose. Washing and sucrose gradient repeated until spores reached 99% purity. Once pure, spores were quantified using a NeuBauer chamber and stored at -80°C until use at a 5×10^9^ spores/mL concentration.

### Spore Extraction Methods

1.5×10^7^ spores pelleted down and resuspended in 50 uL of buffer. Samples with EBB (EBB 8M Urea, 2M Thiourea, 4% w/v SDS, 2% w/v β-mercaptoethanol) were boiled 10 minutes and spun down at 15k rpm for 5 minutes ^23^. Supernatant containing spore extract was saved and spore pellet washed three times with distilled water to remove residual spore extract and buffer. Spores were then retreated with EBB for a total of three extractions, with supernatant saved between each step. USD (8M Urea, 1% w/v SDS, 50 mM DTT, 50 mM Tris-HCl pH 8) samples were treated for 90 minutes at 37°C and pelleted down^30^. Supernatant was saved and spore pellet washed. 1° USD or 1° EBB treated spores were incubated with lysozyme for 37°C overnight.

### Lysozyme Treatment

*C. difficile* spores were treated with lysozyme to analyze the effects of the extraction buffers on the integrity of the spore coat^29^. Spores (1×10^7^ spores) were treated with 1mg/mL lysozyme in 25 mM phosphate buffer (pH 7.4) at 37°C for 2 hours ^10^. Samples were washed with distilled water and analyzed under phase contrast microscopy. 200 spores were analyzed for germination and binned for phase dark, phase grey, or phase bright. The data represents the results of three independent experiments and spore sample preparations.

### Transmission Electron Microscopy and Analysis

For transmission electron microscopy, we followed prior protocols with modifications^31^.Briefly, *C. difficile* spores were fixed overnight with 3% glutaraldehyde, 0.1 M cacodylate buffer (pH 7.2) at 4°C. Samples were centrifuged at 14k rpm for 5 minutes and supernatant discarded. They were then stained with1% osmium tetroxide in 0.05M HEPES buffer (pH 7.4) overnight at 4°C. Treated samples were washed 5x with distilled water. The spores were dehydrated stepwise in 30%, 50%, 70%, 90% for 15 minutes respectively, followed by dehydration with 100% acetone 3 times for 30 minutes at each step. A small amount of acetone is left covering the sample to prevent rehydration. Spore samples were embedded in modified Spurr’s resin (Quetol ERL 4221 resin; EMS; RT 14300) in a Pelco Biowave processor (Ted Pella, Inc.). Initially, 1:1 acetone-resin for 10 min at 200W-with no vacuum, 1:1 acetone-resin for 5 min at 200W-vacuum 20”Hg, followed by 100% resin 4 times at 200W for 5 minutes-vacuum 20”Hg. The resin was removed, and sample pellet was transferred to BEEM conical-tip capsule and filled with 100% fresh modified Spurr’s resin. Sample was left to reach bottom of capsule and subsequently left to polymerize for 48 hours in a 65°C oven followed by 24 hours at room temperature. Ultrathin section ∼100nm (silver-gold color) were obtained using Leica UC7 Ultramicrotome and placed on glow-discharged carbon coated 300-mesh Cu grids. Grids were double lead stained with 2% uranyl acetate for 5 minutes and washed with filter sterilized (0.2uM filter) distilled water followed by 5 minute staining with Reynold’s lead citrate and subsequent washing as described. All ultrathin TEM sections were imaged on a JEOL 1200 EX TEM (JEOL, Ltd.) at 100 kV, and images were recorded on an SIA-15C charge-coupled device (CCD) (Scientific Instruments and Applications) camera at the resolution of 2,721 by 3,233 pixels using MaxImDL software (Diffraction Limited). All equipment used is located at the Texas A&M University Microscopy and Imaging Center Core Facility (RRID: SCR_022128).

ImageJ was used to measure all samples in nanometers. Initially, the spore length and core width was measured three times and the average was calculated. Cortex width was measured once from inner membrane to cortex. Statistical analysis using ROUT to remove outliers and data was analyzed using one way analysis of variance (ANOVA) with Tukey test for multiple comparison. Statistical cutoff for significance with a P-value <0.05 was used.

### Western Blot Analysis

Protein samples were resuspended in 2x SDS-PAGE loading buffer (BioRad) with 5% b-mercaptoethanol and boiled for 10 minutes. Samples were run in a 12% acrylamide SDS-PAGE gels. Proteins were transferred to nitrocellulose membrane and blocked at 4°C overnight in 3% bovine serum albumin (BSA) in Tris-buffer saline (TBS) (pH7.4) with 0.1% TWEEN20 (TBS-T). Western blots probed with 1:10,000 of primary antibody of mouse anti-CdeC anti-CdeM, and a 1:15,000 rabbit anti-SleC in 1% BSA in TBS-T for 1h at room temperature ^31^. SleC was a gift from Dr. Joseph Sorg at Texas A&M University. Membrane was washed 3 times for 5 minutes with TBS-T and incubated in secondary antibody 1:10,000 dilution goat anti-mouse HRP (Sigma) in 1% BSA in TBS-T.

### Antibody Preparation

*E. coli* BL21 (DE3) pRIL was used to overexpress CdeC and CdeM proteins. CdeC and CdeM were respectively overexpressed using pETM11 vector with a C-terminal 6xHIS tag. Plasmids were transformed into BL21 (DE3) pRIL cells and inoculated in 5 mL of Leuria Bertani (LB) broth supplemented with 50 ug/mL chloramphenicol, 10 ug/mL tetracycline, and 50 ug/mL kanamycin in a shaker overnight at 37°C. Overnight culture was used to inoculate a 300 mL flask with the necessary antibiotics at a 1:100 ratio and grown at 37°C until OD600 reached 0.7-0.8. Fresh LB with the appropriate antibiotics was added until flask reached 1L volume. Once OD600 was reached culture was induced with 0.5 mM isopropyl-β-D-1-thiogalactopyranoside (IPTG) for 18 hrs at 21°C. Cultures were pelleted down at 6500 rpm for 10 minutes and stored at -80°C. Pellet was resuspended in 5mL of soluble buffer (50 mM NaH2PO4, 300 mM NaCl, 20 mM imidazole, 1mM phenylmethanesulfonyl fluoride (PMSF), pH=8) and sonicated 15 seconds on, 15 seconds on ice for a total of 6 cycles at 50% amplitude. Lysate was pelleted down at 6500 rpm at 4°C for 30 minutes and subsequently filtered with a 0.22 uM filter.

Purification of recombinant CdeC and CdeM was done using Akta Start. Briefly, filtered soluble protein extract were loaded on HisTrap FF crude column (GE Healthcare), washed with 15 column volumes (CV) of wash buffer (50 mM NaH2PO4, 300 mM NaCl, and 20 mM imidazole, pH=8). Proteins were eluted with 10 CV of elution buffer (50 mM NaH2PO4, 300 mM NaCl, and 250 mM imidazole, pH=8). Samples were eluted as a single peak and subsequent fractions were analyzed by 15% SDS-PAGE and stained with Coomassie G250 to determine purity of fractions. To quantify protein Pierce™ BCA Protein Assay Kit (ThermoFisher Scientific) was used.

5 BALB/c mice, 7-9 weeks old were used for immunization with CdeC and CdeM respectively. A pre-immunized bleed was taken at day -1 from isofluorene anesthetized mice. Blood was incubated for 30 minutes at room temperature to coagulate and then centrifuged at 5000 rpm for 10 minutes at 4°C. Serum was taken and stored in a new tube. At day 0, 20 ug of recombinant protein was emulsified with Freund’s Complete Adjuvant (FCA) in a 1:1 ratio. Mice were anaesthetized and injected with 200 uL of protein emulsified in FCA subcutaneously in the nape. Mice were inoculated as described at day 14 and 28 however using Freund’s Incomplete Adjuvant (FIA). At day 42, mice were anaesthetized with 5% isoflurane and maintained at 1.5% and a cardiac puncture was performed to collect blood. Serum was collected as previously described. Animal protocol was approved by the Comité de Bioética of the Facultad de Ciencias Biológicas at the Universidad Andrés Bello under the approval act code 0035/2018.

## Results and discussions

### Experimental design of spore coat and exosporium extraction

Two chemical methods to extract the spore coat and exosporium proteins have been extensively utilized in *C. difficile* spore-research ^10, 15–23^. However, their extraction efficiency of these two outermost layers remains unclear. To compare treatments, initially as shown in Figure 1, spores were treated with either EBB or USD. EBB treated samples were boiled 10 minutes and then spun down, supernatant containing spore coat and exosporium extracts was saved and remaining spore pellet was washed to remove residual extraction buffer (Fig. 1), a second and third subsequent extraction were performed to remove and identify any residual protein not removed with a single extraction. In the case of USD, spores were incubated at 37°C for 90 minutes and after incubation spun down (Fig. 1); Supernatant was saved, and spore pellet was washed. A second and third subsequent extraction were performed as described. To analyze proteins within the core and cortex, spores treated with USD or EBB, 1°, were incubated with 10 mg/mL of lysozyme overnight. The treatment with lysozyme would allow releasee of proteins within the cortex and core that were not able to be extracted with EBB or USD alone. For all treatments, exosporium/spore coat extracts and treated spores were run through SDS PAGE and immunoblotted for western blot analysis (Fig. 1).

**Figure 1.**
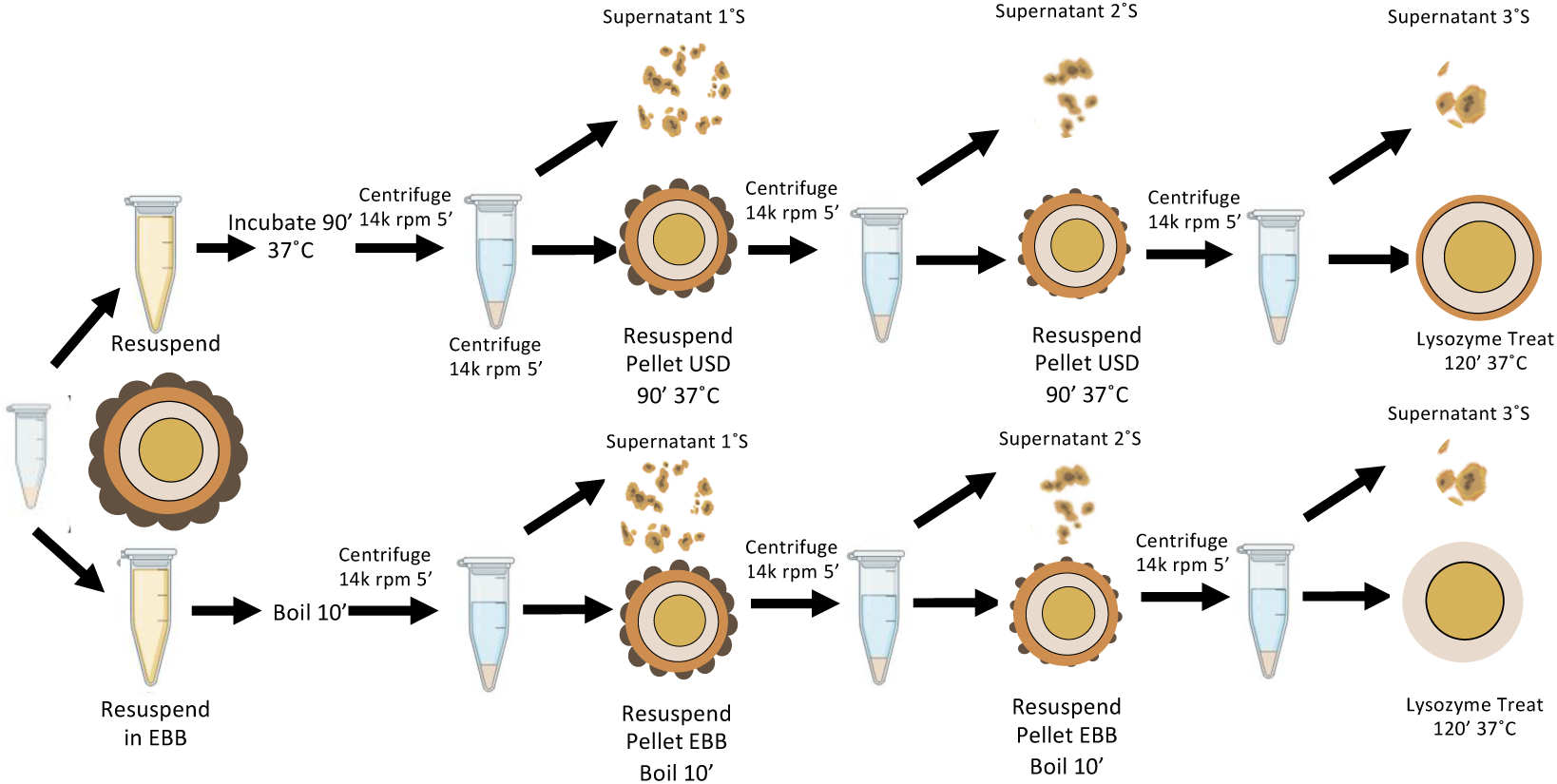
Schematic of extraction treatment. Purified R20291 *C. difficile* spore (5×10^9^ spores/mL) were pelleted down and resuspended in USD or EBB. USD treated spores were incubated at 37°C for 90’ min. EBB treated spores were boiled for 10’ min. Spores were pelleted down for 5’min at 14,000 rpm and supernatant 1°S saved for SDS-PAGE and Western Blot Analysis. Spore pellet (1°) was washed 3x with distilled H2O and centrifuged in between washes at 14,000rpm for 5’min. 1° treated spores were subsequently treated two more times with USD or EBB as described above denoted as 2° and 3°. Supernatant and a portion of spore pellet is saved between each step for analysis. 1° spores were treated with 10mg/mL for 24 hours at 37°C to identify effects of treatments on spore coat integrity.

### Transmission Electron Microscopy of Treated Spores and Measurements

To visualize the efficiency of extraction of the spore surface layers 1° treated spores with USD, and EBB were imaged through TEM. As a controls, untreated and Laemmli-treated spores were also included. As seen in Figure 2, wild type treated spores have electron dense outer layer, the exosporium, the classical electron-dense bumps shown in thick-exosporium spores, and hair-like projections, followed by the underlying spore coat, cortex, and core (Fig. 2A). However, a single EBB- or USD-treatment was able to remove the majority of the spore coat and exosporium layers, removes the majority of the outermost outermost layers, leaving the spore-peptidoglycan cortex exposed (Fig. 2A). Electron micrographs of Laemmli-treated spores had a thin layer of electron dense material in the outermost area of the spore, suggesting that significant spore coat/exosporium residual material remains in the spore surface (Fig. 2A). Altogether, these results demonstrate that, while all three treatment remove the spore coat and exosporium layers, EBB and USD are more efficient.

**Figure 2.**
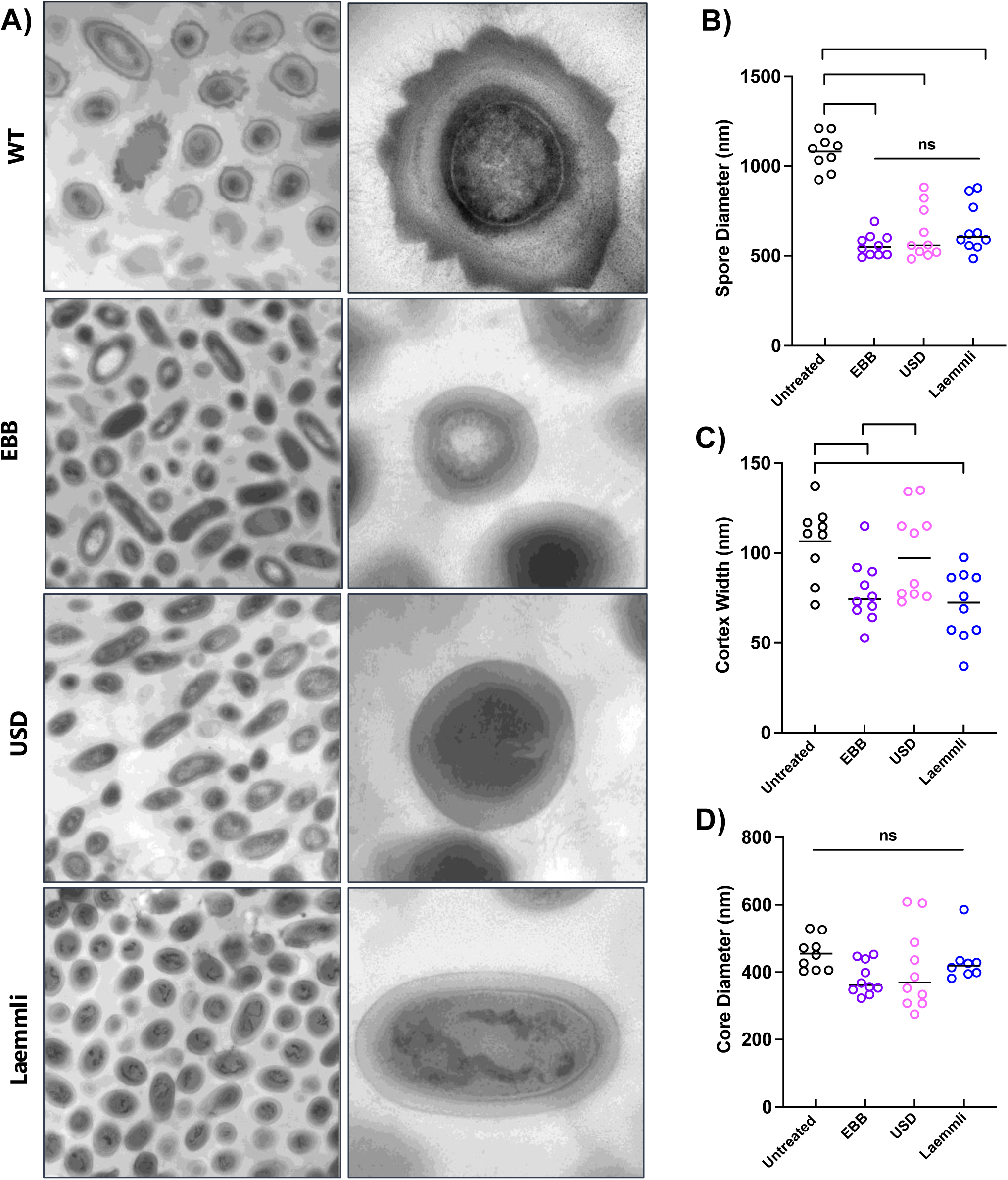
Transmission Electron Microscopy of Treated *C. difficile* spores. A) Transmission Electron Microscopy images of Wild Type, EBB, USD, and Laemmli treated *C. difficile* spores. Measurements were taken using ImageJ of spore diameter, core width, and core diameter (B to D) 9 wild type spores and 10 of each treated spore were measured. Statistical significance was performed using a 1 way ANOVA with Tukey’s multiple comparisons test (not significant-ns, * p < 0.05; ** p < 0.01, *** p<0.001, **** p<0.00001).

To quantitatively assess the impact of these chemical treatments on the ultrastructure of the spore layers, the length of the entire spore, cortex and spore core where measured. The spore length significantly decreased by ∼40 % in spores treated with Laemmli, USD, and EBB when compared to untreated spores (Fig. 2B). *C. difficile* R20291 spores had an average length of ∼ 1,000 nm whereas on average Laemmli-, USD-, and EBB-treated spores were ∼ 600 nm in length (Fig. 2B). Comparing effect of treatment on the spore core indicated a decrease in core diameter in all the treated samples, however this was not statistically significant. (Fig 2C). There was a ∼20% reduction in core diameter in EBB treated spores compared to untreated (Fig. 2C). The average diameter of the core in untreated spores was 455 nm compared to 363 nm in EBB treated spores (Fig. 2C). When comparing the cortex there was significant decrease in cortex between Laemmli and EBB, and Laemmli and USD treated samples (Fig 2C). The decreased cortex width of EBB and Laemmli treated spores could be potentially due to boiling step during extraction leading to significant shrinkage in width, step that is absent in USD-treated spores (Fig. 1). No significant differences where observed in spore core width between untreated and Laemmli-treated spores, and between USD- and EBB-treated spores (Fig. 2D). Collectively, these results demonstrate that while all three treatments (i.e., Laemmli, USD and EBB) remove the spore coat and exosporium layer, those with a boiling step (i.e., Laemmli and EBB) affect the width of the spore cortex, but not spore core.

### Impact of EBB and USD-treatments in spore coat and exosporium extracts

Having demonstrated that all three chemical treatments remove most of the spore surface layers, we thought to correlate the ultrastructure of spores treated with EBB, USD or Laemmli with the efficiency of removal of these spore surface proteins by analyzing their protein-profile by SDS-PAGE. For this we treated the spores with EBB, USD or Laemmli, spun down and the spore-pellet and supernatant electrophoresed in an SDS-PAGE (Fig. 1 and 3A,B). Notably, a spore coat and exosporium EBB extracts from a single EBB-treatment resolved a significantly more protein than Laemmli-extracted spore coat and exosporium proteins (Fig. 1 and 3A). Indeed, comparison of the remaining pellet of EBB- and Laemmli-treated spores shows that EBB-treated spores have less remaining spore coat / exosporium material attached (Fig. 1 and 3A). A second (2°) and third (3°) EBB treatment led to complete removal of the spore coat and exosporium extracts and spore-pellets containing negligible remnants of spore coat and exosporium material (Fig. 1 and 3A). In contrast to EBB, although a first USD-treatment yielded substantial protein levels of spore coat extracts, several high and low-molecular weight protein species remained in USD-spore pellets. Indeed, a second USD-treatment led to significant spore coat and exosporium extracts being removed of USD-treated spores (Fig. 3B). No additional protein was removed after a third USD extraction (Fig. 3B). However, spore pellets from a second and third treatment with USD still retained detectable levels of low molecular mass protein species (∼10 to 15 kDa) (Fig. 3B). Collectively, these results indicate that EBB is more effective in spore coat and exosporium extraction from *C. difficile* spores than USD and Laemmli.

**Figure 3.**
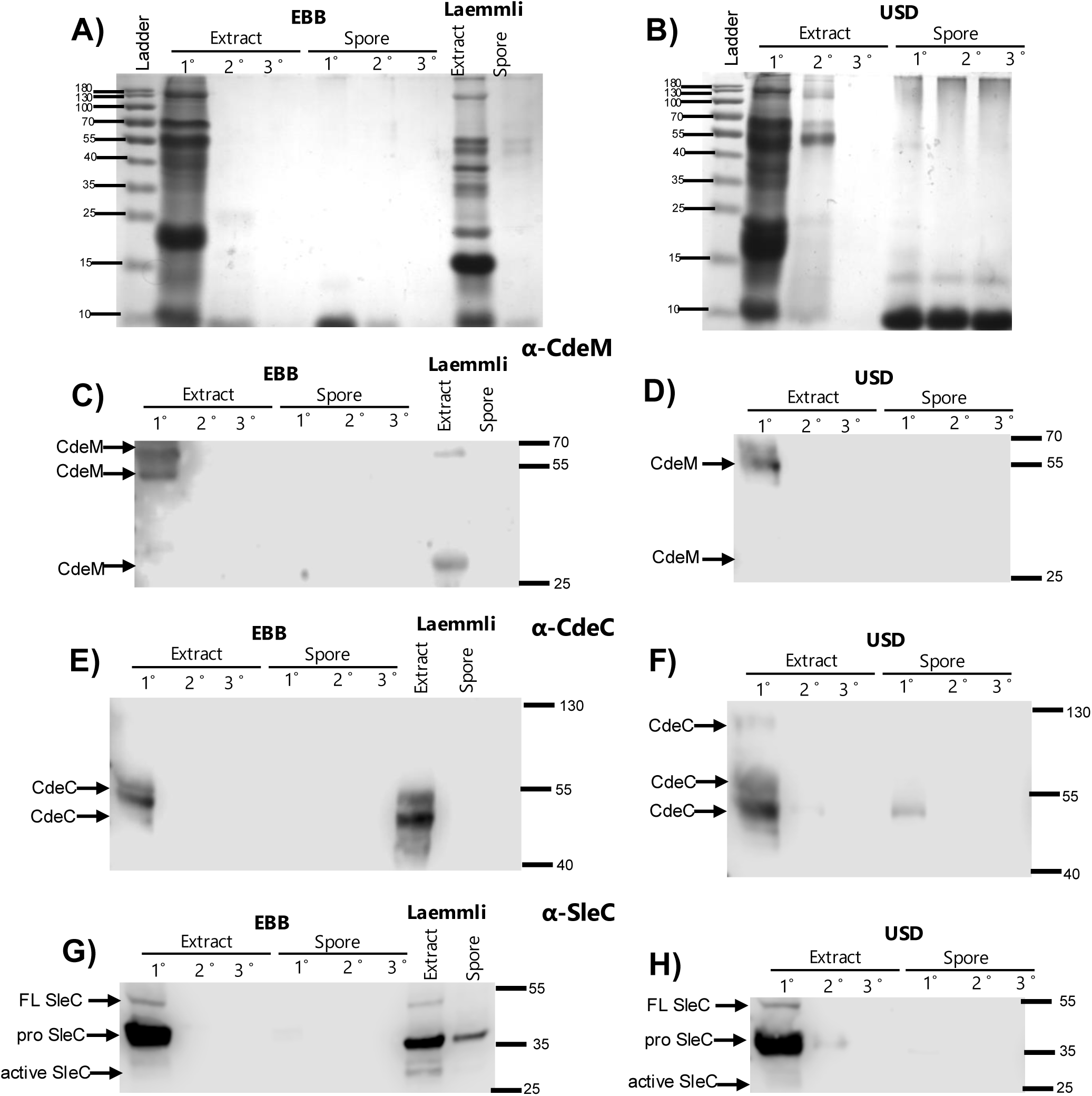
Extraction of Spore Outer layers using EBB and USD. **A)** 15% SDS-PAGE of purified R20291 spores treated with USD, EBB, and 2x Laemmli. Spores were treated as described in Methods and Figure 1. 1°,2°, 3° indicate consecutive extraction treatment of spores, Laemmli 1° indicates one time Laemmli extraction. Western blot analysis of treated samples using: **B)** anti-CdeM antibodies, arrows indicate immunoreactive bands at ∼70 kDa, 55 kDa, and 27 kDa respectively. **C)** anti-CdeC antibodies, arrows indicate immunoreactive bands indicating extraction of CdeC with bands at ∼130 kDa, 55 kDa, and 37 kDa, respectively. **D)** anti-SleC antibodies indicate immunoreactive bands at 55kDa full length SleC, 37 kDa pro-SleC, and 27kDa active SleC. Higher olimeric bands likely oligomerization of the protein.

### Impact of chemical treatments in removal of exosporium and spore coat protein markers

While the above electrophoresed protein profiles correspond to total spore coat and exosporium extracts, the removal efficiency of each of these layers is unclear. Therefore, we utilized antibodies against three proteisn as markers. For the spore coat marker, we selected the cortex-lytic enzyme SleC, known to be located in the interface between the spore cortex ^32^. For exosporium markers, we selected the cysteine-rich and morphogenetic proteins, CdeC and CdeM ^15, 17, 29^; where, CdeM is located on the outermost layer of the exosporium whereas CdeC is found within the exosporium with presumable interactions with the spore coat ^17, 33^.

For this, first, aliquots of the 1^st^ EBB extraction blotted against anti-CdeM shows immunoreactive bands at ∼70 kDa, ∼55 kDa, and ∼ 27 kDa (Fig. 3C). In the case of subsequent EBB extractions (2° and 3°), there were no immunoreactive bands in the extracted supernatant as well as in the spore pellet, indicating that the exosporium layer was completely removed (Fig. 3C). Again, Laemmli-treated spores were used as a control. Supernatant of Laemmli-treated spores containing spore coat and exosporium extracts show the presence of ∼ 70 and 27 kDa immunoreactive species (Fig. 3C), while Laemmli spore-pellet completely lacked CdeM immunoreactive bands (Fig. 3C). Immunoblotting using anti-CdeM antibodies of 1^st^ USD spore extracts display immunoreactive bands at ∼ 70 kDa and 55 kDa, however no 27 kDa immunoreactive band was detectable, and no subsequent CdeM-specific immunoreactive band was observed in subsequent USD spore coat / exosporium extracts (Fig 3D). Moreover, no remnants of CdeM were detected in the USD-treated spore pellet (Fig. 3D). These results demonstrate that all three treatments efficiently remove the exosporium layer but differ in their levels of dissociation of the various CdeM oligomeric species present in the spore coat / exosporium extracts.

Next, to explore whether CdeC, which is an exosporium protein suggested to be at the interface of the spore coat and the exosporium electron dense layer^10, 17^, was removed as efficient as CdeM, immunoblots where analyzed against anti-CdeC antibodies. Results demonstrate that a single treatment of EBB or Laemmli was sufficient to extract all immunodetectable CdeC, with no CdeC remnants in spore pellets (Fig. 3E,F). In the case of USD-treatment, immunoreactive bands were observed at ∼130, 70 and and 55 kDa, however residual ∼55 kDa immunoractive band was detected in USD-treated spores, which was successfully removed after a second treatment with USD (Fig. 3F). These results demonstrate that EBB and Laemmli are more efficient than USD at extracting proteins that are at the interface with the spore coat, such as CdeC.

Finally, the a spore coat / peptidoglycan interface marker protein, SleC, was used to detect efficient extraction of spore surface layers as well as cortex proteins^31^. During sporulation SleC is processed and appears as three immunoreactive bands in spore coat / exosporium extracts, where full length ∼55 kDa, pro-SleC ∼37 kDa, and active SleC ∼27 kDa^32^. An initial EBB extraction displayed immunoreactive bands for all three versions of SleC at the expected ∼55 kDa, ∼37 kDa, and ∼27 kDa, while EBB-treated spore pellets had a very faint band at ∼37 kDa (Fig 3G), suggesting the presence of remnants at the interface of the spore peptidoglycan / spore coat layers or remnants of the spore coats themselves. A second EBB-treatment was sufficient to completely remove SleC remnants (Fig. 3G). Notably, Laemmli gave consistent immunoreactive bands for all forms of SleC but failed to completely remove SleC as seen with a strong immunoreactive band within the treated spore at ∼37 kDa (Fig 3G, 3H). In the case of USD, an initial USD-treatment extracted the majority of SleC, with a faint 37 kDa immunoreactive present in the spore pellets, which was also present upon treating USD-treated spores with a second USD-treatment and completely removed SleC from the spore pellet fraction (Fig 3H). These observations indicate that a single EBB-treatment is sufficient to remove the majority of SleC from the spore coat, and very likely other spore coat proteins.

### Impact of EBB and USD in lysozyme-triggered germination

The spore coat layer is known to act as an impermeable barrier to molecules > 5 kDa, such as enzymes, including proteinase K and lysozyme ^30, 34, 35^. Moreover, the spore peptidoglycan cortex in decoated spores is susceptible to lysozyme degradation, artificially triggering spore germination and outgrowth ^30^. We reasoned that depending on the efficiency with which EBB and USD removed the spore coat, EBB and USD-treated spores will have different levels of susceptibility to lysozyme-germination. Phase contrast analysis of untreated, EBB-, or USD-treated spores reveals that untreated and USD-treated spores are phase bright (Fig. 4A,B), consistent with them remaining dormant. By contrast, the majority (∼90 %) of EBB-treated spores become phase grey (Fig. 4A,B), which could be attributed to the boiling step of *C. difficile* spores, which is known to release spore core DPA contents in *C. difficile* and other spore-formers ^17, 36^.

**Figure 4.**
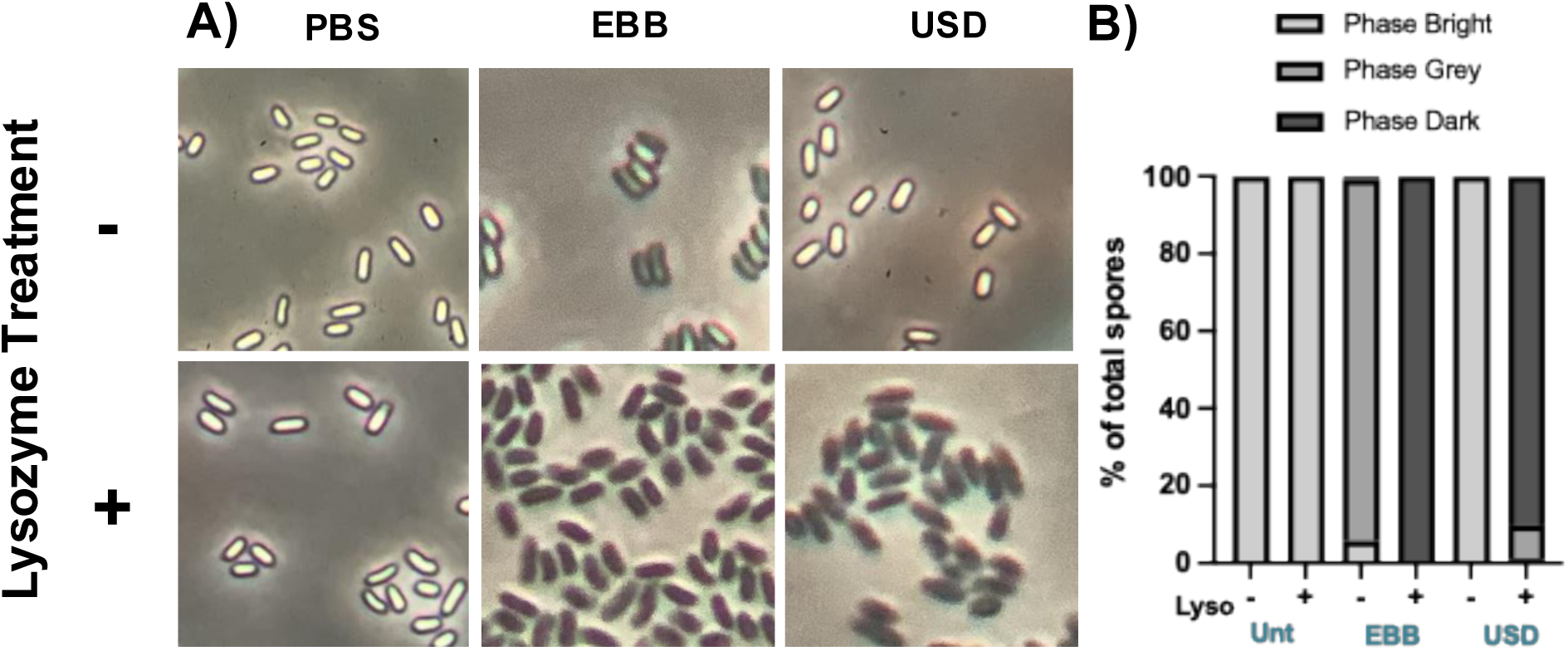
Effect of extraction on spore coat permeability to lysozyme. Effects of USD and EBB treatment on spore coat integrity and lysozyme resistance. R20291 *C. difficile* spores were treated for 90 min at 37°C incubator with USD, washed and imaged under phase contrast. EBB treated spores were boiled for 10 min, centrifuged for 5 min at 14,000 rpm and washed. Spores were treated with 10 mg/mL lysozyme for 24 hours at 37°C. Effects on spore coat integrity and degradation of peptidoglycan indicated by phase dark spores under phase contrast. B) Lysozyme treated spores were quantified and percent phase bright, grey, and dark were graphed.

To assess effects of USD and EBB treatment on spore coat permeability and extraction efficiency, spores were treated with lysozyme (10 mg/mL) overnight at 37 °C for 24 hours ^17, 23^. Untreated spores appeared phase bright, which is indicative that lysozyme was unable to trigger germination. By contrast, most of the EBB and USD treated spores became phase dark which indicates that lysozyme degraded the cortex, leading to spore core hydration (Fig. 4A,B). However, a subfraction of USD-treated spores subsequently treated with lysozyme appeared phase grey. This suggests that unlike phase dark spores, phase grey spores either partially released their spore core DPA and/or exhibit partial spore core rehydration. Alternatively, it is likely that phase grey spores retain remnants of the spore coat which impedes lysozyme-germination and subsequent spore core hydration. Collectively, these observations demonstrate that EBB-treatment provides a homogenous population of decoated spores that can be utilized in downstream applications.

### Impact of lysozyme digestion in the protein profile of EBB- and USD-treated spores

The results showing that EBB and USD remove most of the spore coat and that these decoated spores are readily digested with lysozyme led us to speculate whether these steps could be utilized to pop-open the spore core for electrophoresis. As observed, resuspending EBB-, USD- or Laemmli-treated spores in Laemmli sample buffer followed by 5 min incubation at 90 °C prior to electrophoresing them in an SDS-PAGE gel (Fig. 5), did not release the proteins contained in the spore peptidoglycan cortex and in the spore core, demonstrating the resilience of decoated spores to harsh treatments. However, upon incubating EBB- or USD-decoated spores with lysozyme for 24 hrs at 37 °C and subsequently electrophoresed, we observed an increase in the levels of proteins being electrophoresed (Fig. 5), which improved further when incorporating Laemmli buffer and a 5 min incubation at 90 °C prior to electrophoresis (Fig. 5). Notably, despite Laemmli buffer being able to remove the spore coats (Fig. 2 and 3), Laemmli-decoated spores subsequently treated with lysozyme did not release spore core content material. In summary, these results evidence that electrophoresing lysozyme-treated decoated spores provides a method to release the spore peptidoglycan cortex and spore core proteins.

**Figure 5.**
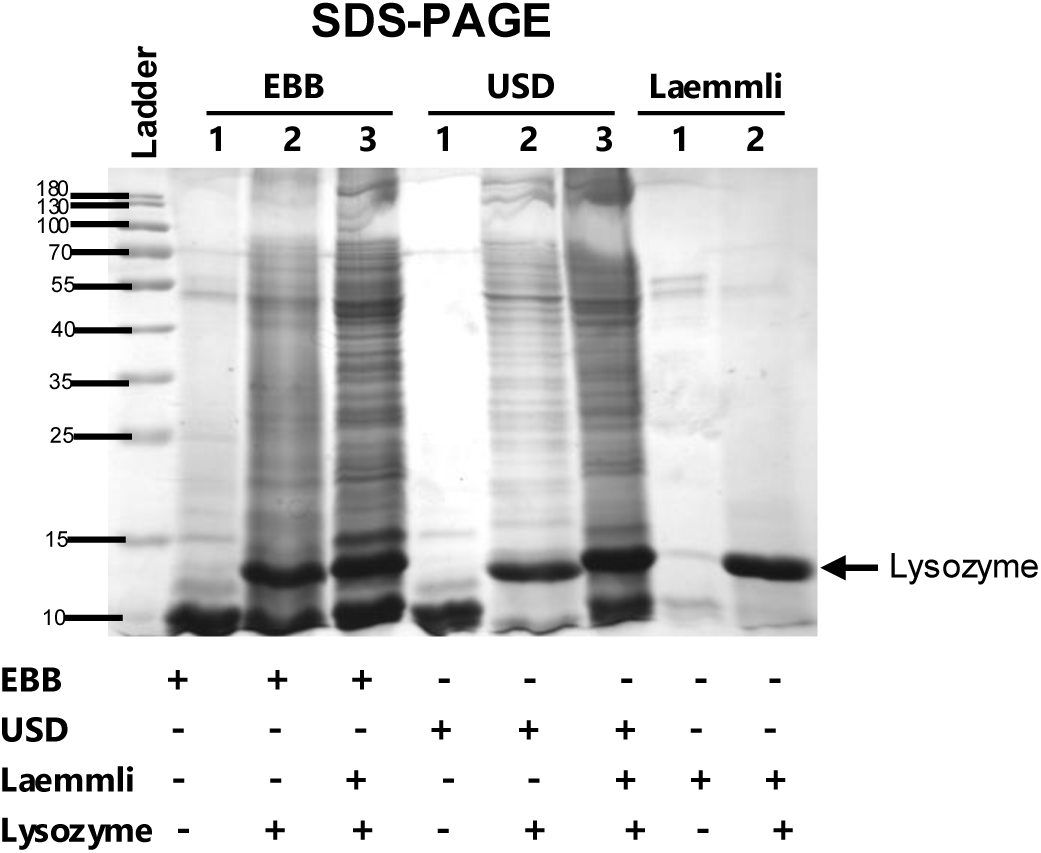
Lysozyme treatment of decoated-spores. 15% SDS-PAGE of purified R20291 spores treated with USD, EBB, and 2x Laemmli. Spores were treated as described in Methods and Figure 1. 1° and 2° spores treated with 10mg/mL lysozyme, 1°spores were not treated with lysozyme after initial USD/EBB treatment. 1°Laemmli treated spores with and without lysozyme also shown. ∼14kDa band on SDS corresponding to lysozyme indicated.

## Conclusions

This work provides evidence that EBB-decoating treatment is more efficient than USD in removing the spore coat and exosporium layers. Additionally, results also demonstrate that EBB-treated spores can be digested with lysozyme to release the spore peptidoglycan cortex and spore core proteins. These techniques can be adapted in methods seeking to fractionate the different layers of the spore (i.e., exosporium, spore coat, spore peptidoglycan cortex, and spore core).

## Acknowledgments

This project was supported by a grant from the National Institute of Allergy and Infectious Diseases NIAID Grant R01AI177842 to D.P-S.

## References

1. Evans CT, Safdar N. Current Trends in the Epidemiology and Outcomes of *Clostridium difficile* Infection. Clin Infect Dis. 2015;60 Suppl 2:S66–71. doi: 10.1093/cid/civ140. PubMed PMID: 25922403.

2. Ramsay I, Brown NM, Enoch DA. Recent Progress for the Effective Prevention and Treatment of Recurrent *Clostridium difficile* Infection. Infect Dis (Auckl). 2018;11:1178633718758023. Epub 2018/03/15. doi: 10.1177/1178633718758023. PubMed PMID: 29535530; PMCID: PMC5844436.

3. Stevens VW, Nelson RE, Schwab-Daugherty EM, Khader K, Jones MM, Brown KA, Greene T, Croft LD, Neuhauser M, Glassman P, Goetz MB, Samore MH, Rubin MA. Comparative Effectiveness of Vancomycin and Metronidazole for the Prevention of Recurrence and Death in Patients With *Clostridium difficile* Infection. JAMA Intern Med. 2017;177(4):546–53. doi: 10.1001/jamainternmed.2016.9045. PubMed PMID: 28166328.

4. Rupnik M, Wilcox MH, Gerding DN. *Clostridium difficile* infection: new developments in epidemiology and pathogenesis. Nat Rev Microbiol. 2009;7(7):526–36. Epub 2009/06/17. doi: 10.1038/nrmicro2164. PubMed PMID: 19528959.

5. Deakin LJ, Clare S, Fagan RP, Dawson LF, Pickard DJ, West MR, Wren BW, Fairweather NF, Dougan G, Lawley TD. The *Clostridium difficile spo0A* gene is a persistence and transmission factor. Infect Immun. 2012;80(8):2704–11. Epub 2012/05/23. doi: 10.1128/IAI.00147-12. PubMed PMID: 22615253; PMCID: PMC3434595.

6. Paredes-Sabja D, Cid-Rojas F, Pizarro-Guajardo M. Assembly of the exosporium layer in Clostridioides difficile spores. Curr Opin Microbiol. 2022;67:102137. Epub 2022/02/20. doi: 10.1016/j.mib.2022.01.008. PubMed PMID: 35182899.

7. Paredes-Sabja D, Shen A, Sorg JA. *Clostridium difficile* spore biology: sporulation, germination, and spore structural proteins. Trends Microbiol. 2014;22(7):406–16. Epub 2014/05/13. doi: 10.1016/j.tim.2014.04.003. PubMed PMID: 24814671; PMCID: PMC4098856.

8. Castro-Córdova P, Otto-Medina M, Montes-Bravo N, Brito-Silva C, Lacy DB, Paredes-Sabja D. Redistribution of the Novel Clostridioides difficile Spore Adherence Receptor E-Cadherin by TcdA and TcdB Increases Spore Binding to Adherens Junctions. Infect Immun. 2023;91(1):e0047622. Epub 20221130. doi: 10.1128/iai.00476-22. PubMed PMID: 36448839; PMCID: PMC9872679.

9. Castro-Córdova P, Mora-Uribe P, Reyes-Ramírez R, Cofré-Araneda G, Orozco-Aguilar J, Brito-Silva C, Mendoza-León MJ, Kuehne SA, Minton NP, Pizarro-Guajardo M, Paredes-Sabja D. Entry of spores into intestinal epithelial cells contributes to recurrence of *Clostridioides difficile* infection. Nat Commun. 2021;12(1):1140. Epub 2021/02/18. doi: 10.1038/s41467-021-21355-5. PubMed PMID: 33602902.

10. Calderon-Romero P, Castro-Cordova P, Reyes-Ramirez R, Milano-Cespedes M, Guerrero-Araya E, Pizarro-Guajardo M, Olguin-Araneda V, Gil F, Paredes-Sabja D. *Clostridium difficile* exosporium cysteine-rich proteins are essential for the morphogenesis of the exosporium layer, spore resistance, and affect *C. difficile* pathogenesis. PLoS Pathog. 2018;14(8):e1007199. Epub 2018/08/09. doi: 10.1371/journal.ppat.1007199. PubMed PMID: 30089172.

11. Mora-Uribe P, Miranda-Cardenas C, Castro-Cordova P, Gil F, Calderon I, Fuentes JA, Rodas PI, Banawas S, Sarker MR, Paredes-Sabja D. Characterization of the Adherence of *Clostridium difficile* Spores: The Integrity of the Outermost Layer Affects Adherence Properties of Spores of the Epidemic Strain R20291 to Components of the Intestinal Mucosa. Front Cell Infect Microbiol. 2016;6:99. Epub 2016/10/08. doi: 10.3389/fcimb.2016.00099. PubMed PMID: 27713865; PMCID: PMC5031699.

12. Paredes-Sabja D, Sarker MR. Adherence of *Clostridium difficile* spores to Caco-2 cells in culture. J Med Microbiol. 2012;61(Pt 9):1208–18. Epub 2012/05/19. doi: 10.1099/jmm.0.043687-0. PubMed PMID: 22595914.

13. Pizarro-Guajardo M, Calderon-Romero P, Castro-Cordova P, Mora-Uribe P, Paredes-Sabja D. Ultrastructural Variability of the Exosporium Layer of *Clostridium difficile* Spores. Appl Environ Microbiol. 2016;82(7):2202–9. Epub 2016/02/07. doi: 10.1128/AEM.03410-15. PubMed PMID: 26850296; PMCID: PMC4807528.

14. Pizarro-Guajardo M, Calderón-Romero P, Paredes-Sabja D. Ultrastructure Variability of the Exosporium Layer of Clostridium difficile Spores from Sporulating Cultures and Biofilms. Appl Environ Microbiol. 2016;82(19):5892–8. Epub 20160916. doi: 10.1128/aem.01463-16. PubMed PMID: 27474709; PMCID: PMC5038037.

15. Diaz-Gonzalez F, Milano M, Olguin-Araneda V, Pizarro-Cerda J, Castro-Cordova P, Tzeng SC, Maier CS, Sarker MR, Paredes-Sabja D. Protein composition of the outermost exosporium-like layer of *Clostridium difficile* 630 spores. J Proteomics. 2015;123:1–13. Epub 2015/04/08. doi: 10.1016/j.jprot.2015.03.035. PubMed PMID: 25849250.

16. Abhyankar W, Hossain AH, Djajasaputra A, Permpoonpattana P, Ter Beek A, Dekker HL, Cutting SM, Brul S, de Koning LJ, de Koster CG. In pursuit of protein targets: proteomic characterization of bacterial spore outer layers. J Proteome Res. 2013;12(10):4507–21. Epub 20130923. doi: 10.1021/pr4005629. PubMed PMID: 23998435.

17. Barra-Carrasco J, Olguín-Araneda V, Plaza-Garrido A, Miranda-Cárdenas C, Cofré-Araneda G, Pizarro-Guajardo M, Sarker MR, Paredes-Sabja D. The Clostridium difficile exosporium cysteine (CdeC)-rich protein is required for exosporium morphogenesis and coat assembly. J Bacteriol. 2013;195(17):3863–75. Epub 20130621. doi: 10.1128/jb.00369-13. PubMed PMID: 23794627; PMCID: PMC3754587.

18. Pishdadian K, Fimlaid KA, Shen A. SpoIIID-mediated regulation of σK function during *Clostridium difficile* sporulation. Mol Microbiol. 2015;95(2):189–208. Epub 2014/12/19. doi: 10.1111/mmi.12856. PubMed PMID: 25393584; PMCID: PMC4377281.

19. Fimlaid KA, Bond JP, Schutz KC, Putnam EE, Leung JM, Lawley TD, Shen A. Global analysis of the sporulation pathway of *Clostridium difficile*. PLoS Genet. 2013;9(8):e1003660. Epub 2013/08/08. doi: 10.1371/journal.pgen.1003660. PubMed PMID: 23950727; PMCID: PMC3738446.

20. Putnam EE, Nock AM, Lawley TD, Shen A. SpoIVA and SipL are *Clostridium difficile* spore morphogenetic proteins. J Bacteriol. 2013;195(6):1214–25. Epub 20130104. doi: 10.1128/jb.02181-12. PubMed PMID: 23292781; PMCID: PMC3592010.

21. Abhyankar WR, Zheng L, Brul S, de Koster CG, de Koning LJ. Vegetative Cell and Spore Proteomes of Clostridioides difficile Show Finite Differences and Reveal Potential Protein Markers. J Proteome Res. 2019;18(11):3967–76. Epub 20191014. doi: 10.1021/acs.jproteome.9b00413. PubMed PMID: 31557040; PMCID: PMC6832669.

22. Swarge BN, Roseboom W, Zheng L, Abhyankar WR, Brul S, de Koster CG, de Koning LJ. “One-Pot” Sample Processing Method for Proteome-Wide Analysis of Microbial Cells and Spores. Proteomics Clin Appl. 2018;12(5):e1700169. Epub 20180416. doi: 10.1002/prca.201700169. PubMed PMID: 29484825; PMCID: PMC6174930.

23. Lawley TD, Croucher NJ, Yu L, Clare S, Sebaihia M, Goulding D, Pickard DJ, Parkhill J, Choudhary J, Dougan G. Proteomic and genomic characterization of highly infectious Clostridium difficile 630 spores. J Bacteriol. 2009;191(17):5377–86. Epub 20090619. doi: 10.1128/jb.00597-09. PubMed PMID: 19542279; PMCID: PMC2725610.

24. Bennion BJ, Daggett V. The molecular basis for the chemical denaturation of proteins by urea. Proc Natl Acad Sci U S A. 2003;100(9):5142–7. Epub 20030417. doi: 10.1073/pnas.0930122100. PubMed PMID: 12702764; PMCID: PMC154312.

25. Romero-Rodríguez A, Troncoso-Cotal S, Guerrero-Araya E, Paredes-Sabja D. The cysteine-rich exosporium morphogenetic protein, CdeC, exhibits self-assembly properties that lead to organized inclusion bodies in Escherichia coli. mSphere. 2020;5 (6):e01065–20. doi: DOI: 10.1128/mSphere.01065-20.

26. Barra-Carrasco J, Olguin-Araneda V, Plaza-Garrido A, Miranda-Cardenas C, Cofre-Araneda G, Pizarro-Guajardo M, Sarker MR, Paredes-Sabja D. The *Clostridium difficile* exosporium cysteine (CdeC)-rich protein is required for exosporium morphogenesis and coat assembly. J Bacteriol. 2013;195(17):3863–75. Epub 2013/06/25. doi: 10.1128/JB.00369-13. PubMed PMID: 23794627; PMCID: PMC3754587.

27. Francis MB, Allen CA, Sorg JA. Spore Cortex Hydrolysis Precedes Dipicolinic Acid Release during Clostridium difficile Spore Germination. J Bacteriol. 2015;197(14):2276–83. Epub 20150427. doi: 10.1128/jb.02575-14. PubMed PMID: 25917906; PMCID: PMC4524186.

28. Paredes-Sabja D, Setlow P, Sarker MR. SleC is essential for cortex peptidoglycan hydrolysis during germination of spores of the pathogenic bacterium *Clostridium perfringens*. J Bacteriol. 2009;191(8):2711–20. Epub 2009/02/13. doi: 10.1128/JB.01832-08. PubMed PMID: 19218389; PMCID: PMC2668406.

29. Calderón-Romero P, Castro-Córdova P, Reyes-Ramírez R, Milano-Céspedes M, Guerrero-Araya E, Pizarro-Guajardo M, Olguín-Araneda V, Gil F, Paredes-Sabja D. Clostridium difficile exosporium cysteine-rich proteins are essential for the morphogenesis of the exosporium layer, spore resistance, and affect C. difficile pathogenesis. PLOS Pathogens. 2018;14(8):e1007199. doi: 10.1371/journal.ppat.1007199.

30. Escobar-Cortes K, Barra-Carrasco J, Paredes-Sabja D. Proteases and sonication specifically remove the exosporium layer of spores of *Clostridium difficile* strain 630. J Microbiol Methods. 2013;93(1):25–31. Epub 2013/02/07. doi: 10.1016/j.mimet.2013.01.016. PubMed PMID: 23384826.

31. Baloh M, Nerber HN, Sorg JA. Imaging Clostridioides difficile Spore Germination and Germination Proteins. J Bacteriol. 2022;204(7):e0021022. Epub 20220628. doi: 10.1128/jb.00210-22. PubMed PMID: 35762766; PMCID: PMC9295549.

32. Gutelius D, Hokeness K, Logan SM, Reid CW. Functional analysis of SleC from Clostridium difficile: an essential lytic transglycosylase involved in spore germination. Microbiology (Reading). 2014;160(Pt 1):209–16. Epub 20131018. doi: 10.1099/mic.0.072454-0. PubMed PMID: 24140647; PMCID: PMC3917228.

33. Díaz-González F, Milano M, Olguin-Araneda V, Pizarro-Cerda J, Castro-Córdova P, Tzeng SC, Maier CS, Sarker MR, Paredes-Sabja D. Protein composition of the outermost exosporium-like layer of Clostridium difficile 630 spores. J Proteomics. 2015;123:1–13. Epub 20150404. doi: 10.1016/j.jprot.2015.03.035. PubMed PMID: 25849250; PMCID: PMC6764588.

34. Alves Feliciano C, Douché T, Giai Gianetto Q, Matondo M, Martin-Verstraete I, Dupuy B. CotL, a new morphogenetic spore coat protein of Clostridium difficile. Environmental Microbiology. 2019;21(3):984–1003. doi: 10.1111/1462-2920.14505.

